# Heterologous production of cyanophycin with *Tatumella morbirosei* cyanophycin synthetase

**DOI:** 10.1101/2023.01.09.523336

**Authors:** Kyle Swain, Itai Sharon, Wyatt Blackson, Stefan Tekel, T. Martin Schmeing, David R. Nielsen, Brent L. Nannenga

**Affiliations:** Chemical Engineering, School for Engineering of Matter, Transport and Energy, Arizona State University, Tempe, AZ 85287 USA; Center for Applied Structural Discovery, The Biodesign Institute, Arizona State University, Tempe, AZ, USA; Department of Biochemistry and Centre de recherche en biologie structurale, McGill University, Montréal, Quebec, Canada

## Abstract

Microbial production of biopolymers represents a promising, sustainable alternative to current approaches for plastic production. Cyanophycin synthetase 1 (CphA1) produces cyanophycin - an attractive biopolymer consisting of a poly-L-aspartic acid backbone decorated with L-arginine side groups. In this work, a series of CphA1 enzymes from different bacteria were screened for heterologous cyanophycin production in engineered *Escherichia coli*, from which it was found that CphA1 from *Tatumella morbirosei* (*Tm*CphA1) was especially productive. *Tm*CphA1 was capable of supporting up to ~2-fold greater yields of insoluble cyanophycin than any other tested CphA1 enzymes, including 10.8-times more than CphA1 from *Synechocystis* sp. PCC6308. Finally, using a bench-scale bioreactor, cyanophycin production by *Tm*CphA1-expressing *E. coli* reached up to 1.9 g per liter of culture by 48 h.

## Introduction

Petroleum-based plastics present significant challenges to the environment due to their synthesis from non-renewable hydrocarbon resources and prolonged degradation time under native conditions [1]. One solution to this problem is to produce biodegradable and renewable polymers from biological sources. Cyanophycin synthetase 1 (CphA1) is a nonribosomal peptide synthetase (unrelated to the family of modular nonribosomal peptide synthetases) that synthesizes cyanophycin, a biopolymer composed of poly-L-aspartic acid and poly-L-arginine subunits [2, 3]. Natively, cyanophycin is produced by species in many bacterial classes as a storage molecule for both nitrogen and carbon, and can later be hydrolyzed into dipeptides by a second enzyme, cyanophycinase (CphB) [4–6]. When isolated, cyanophycin can be separated into insoluble and soluble fractions, which refers to the solubility of the polymer at neutral pH in aqueous solution. Depending on the specific CphA1 enzyme and expression conditions used, lysine can also be incorporated into cyanophycin in place of arginine, and the soluble fraction generally contains a higher ratio of lysine relative to the insoluble fraction [7, 8]. While cyanophycin continues to attract interest as a renewable biopolymer [9], insoluble cyanophycin is particularly attractive for downstream applications due to the significantly reduced processing requirements for purification and higher levels of compositional homogeneity [10].

A prominent challenge for bioplastics production remains the high cost and low yield of most current biopolymers [11]. Therefore, it is of great interest to identify, engineer, and optimize new platforms with enhanced productivity. In this work, we have characterized cyanophycin production in engineered *E. coli* using CphA1 from *T. morbirosei* DSM23827, a Gram-negative rod-shaped bacterium [3, 12]. We found *Tm*CphA1 to support enhanced yields relative to other, more commonly studied CphA1 enzymes. *Tm*CphA is a ~400 kDa tetramer that previously been shown to be functionally expressed in *E. coli* and whose structure was recently determined to 3.1Å by X-ray crystallography [3]. Like other CphA1 enzymes, *Tm*CphA1 has three functional domains that each play an important role in the production of cyanophycin. The G domain uses ATP to catalyze the addition of aspartic acid to the end of the growing cyanophycin chain [2, 13]. The role of the M domain (often compared with Mur ligases) is to add arginine to the sidechain of the recently-added aspartic acid residue, again with the use of ATP [14]. The N domain is thought to use salt bridges and hydrogen bonding to loosely anchor the cyanophycin molecule in place, supporting a “flip-flop” mechanism where the polymer rebounds from the G domain to the M domain and back, allowing individual subunits to be alternately attached to the growing polymer chain [3, 15]. The N domain of most CphA1 enzymes has also recently been shown to generate primers, which are used to initiate cyanophycin synthesis *in vivo*, though *Tm*CphA1 is among the minority of CphA1 enzymes whose N domain lacks this active site [16]. Here we demonstrate that cyanophycin produced by *Tm*CphA1 in *E. coli* is similar in composition to previous systems and, when expressed in a bench-scale bioreactor, enabled the production 1.8 g of insoluble cyanophycin per liter of culture.

## Results and Discussion

Production of cyanophycin in *E. coli* has been demonstrated for several CphA1 enzymes [10, 11, 17]; however, heterologous expression and purification of the stable enzyme in *E. coli* notably remains difficult for many CphA1s [3]. This suggests that, although the synthetase is functional, it is not necessarily stable when expressed in *E. coli*. For example, cyanophycin production by CphA1 from *Synechocystis* sp. PCC6308 (*6308*CphA1) is well-characterized and often serves as a benchmark for heterologous cyanophycin production [18–21], yet the *6308*CphA1 enzyme itself does not robustly express and purify from *E. coli*. Recently, other CphA1 enzymes that do express well in *E. coli* and purify as soluble and stable samples were identified and used for structure determination and functional analysis by two authors of this work [3] and by others [22]. We hypothesized that CphA1 enzymes with robust and stable heterologous expression in *E. coli* would in turn support enhanced cyanophycin production within that host. Accordingly, production of cyanophycin by *E. coli* expressing three different stable enzymes - CphA1 from *Acinetobacter baylyi* DSM587 (*Ab*CphA1), *Synechocystis* sp. UTEX2470 (*Su*CphA1), and *Tm*CphA1 - was characterized and compared with that of *6308*CphA1 in order to identify potential improvements in cyanophycin production resulting directly from the use of a more stable enzyme.

Flask scale (25 mL) cultures were first carried out for *E. coli* expressing each CphA1 enzyme to determine and compare levels of soluble, insoluble, and total cyanophycin production (Fig. 1). While the overall cyanophycin yields of *6308*CphA1, *Ab*CphA1, and *6308*CphA1were similar, *Ab*CphA1 and *Su*CphA1 produced 3.8- and 3.1-times more insoluble cyanophycin than *6308*CphA1, respectively. However, the greatest cyanophycin production was achieved using *Tm*CphA1 which, relative to *6308*CphA1, resulted in 10.8-times more insoluble cyanophycin and 2.6-times more total cyanophycin. Even when compared with *Ab*CphA1, the next best producer of insoluble cyanophycin, 2.9-times more insoluble cyanophycin was produced using *Tm*CphA1. Interestingly, *Tm*CphA1 and *Ab*CphA1 both lack the hydrolytic active site present in the N domains of *6308*CphA1 and *Su*CphA1 [23]. This domain has been shown [23] to increase cyanophycin yield at an earlier time point (20 h) than was assessed here (48 h), but the superior yields obtained for *Tm*CphA1 nevertheless indicate that indicates this “head start” imparted by the N domain hydrolytic site can be overcome in time.

**Figure 1.**
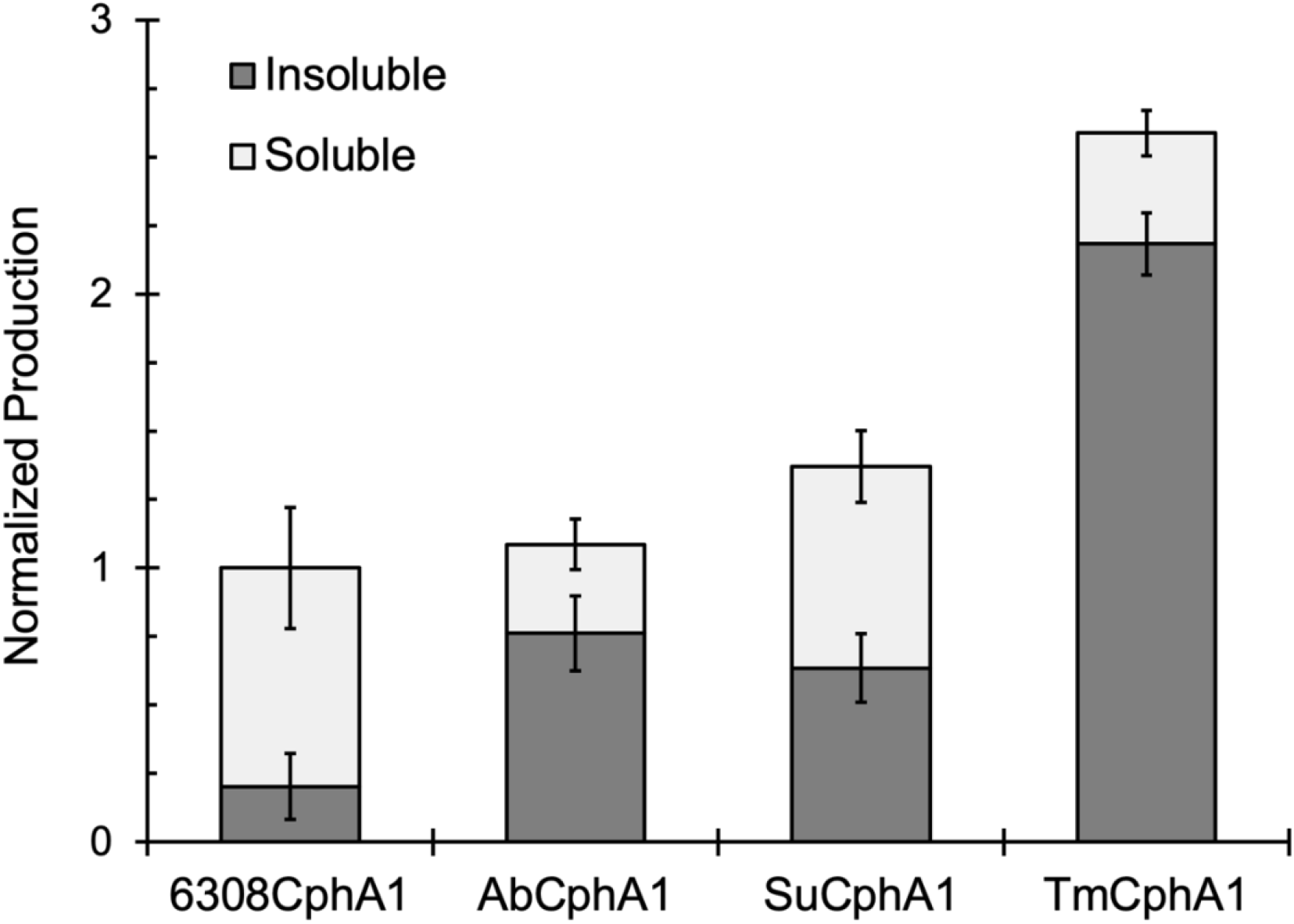
Comparison of cyanophycin production by *E. coli* strains expressing different CphA1 enzymes. *Tm*CphA1 produced more total cyanophycin, especially insoluble cyanophycin, relative to all other homologs tested. Experiments were conducted in 25 mL shake flasks in triplicate and normalized relative to the total cyanophycin produced by *6308*CphA1. Error bars represent the standard deviation.

Due to its enhanced performance, further studies on cyanophycin production by *Tm*CphA1 were next conducted. *Tm*CphA1 expressed in this system was confirmed by size exclusion chromatography (SEC) to be a tetramer, as previously shown [3, 16] (Fig. 2A), and the functionality of purified *Tm*CphA1 was confirmed by the presence of polymer in SDS-PAGE analysis of an *in vitro* assay (Fig 2B). Cyanophycin has a natural size range from 20-100 kDa, with heterologous production in *E. coli* usually generating nearer to 20-40 kDa [24]. Here, cyanophycin produced *in vitro* by purified *Tm*CphA1 showed a size distribution of between 20-30 kDa, similar to what has been shown previously for *Tm*CphA1 [3] and other CphA1 enzymes [15, 25–28].

**Figure 2.**
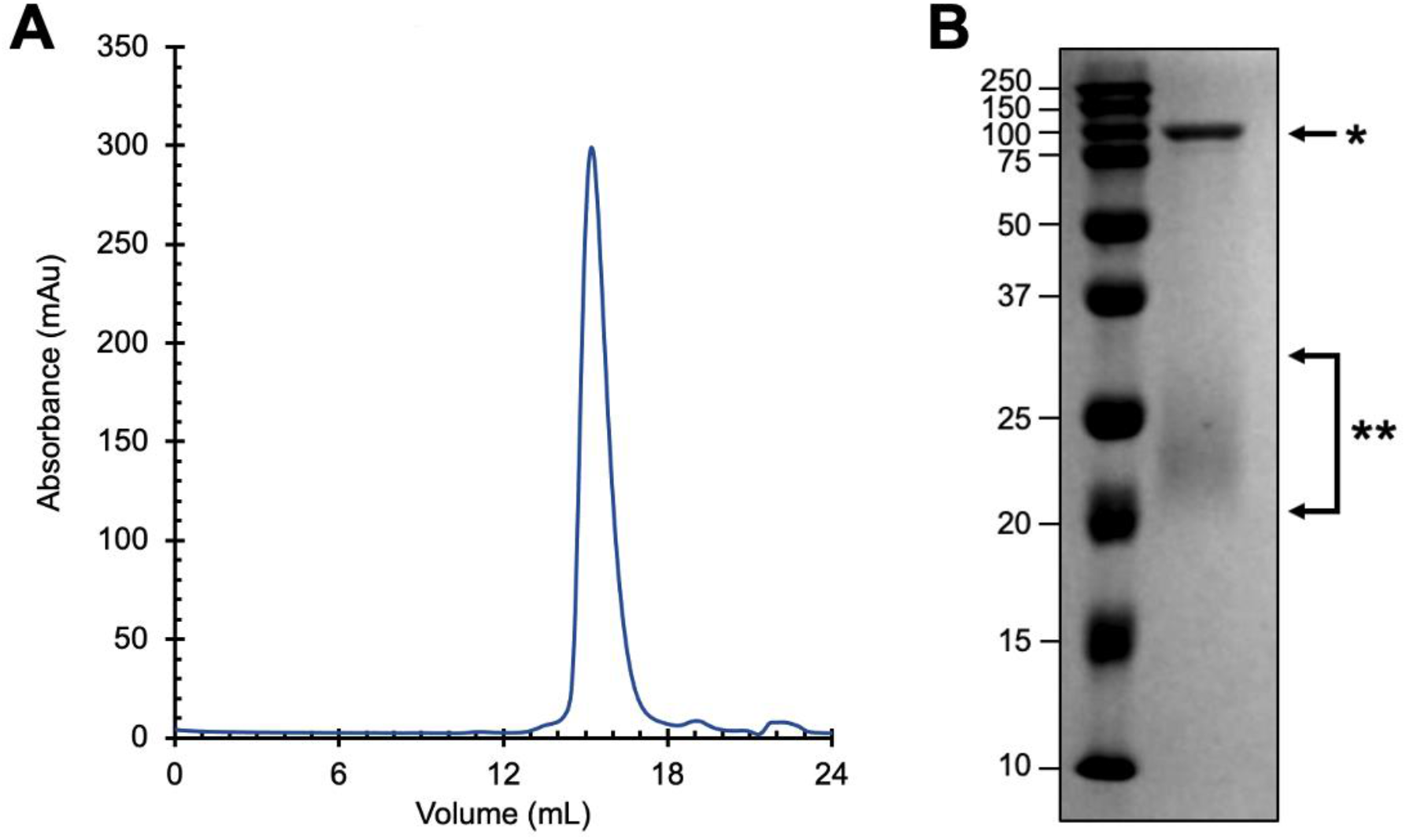
Functional analysis of purified *Tm*CphA1. (A) The SEC chromatograph confirms that *Tm*CphA1 is expressed as a stable homotetramer in *E. coli* with an approximate molecular weight of 400 kDa. (B) SDS-PAGE analysis of purified *Tm*CphA1 (*) shows that the enzyme is active and produces cyanophycin (**) *in vitro* in the presence of ATP, L-aspartic acid, L-arginine, and primer.

The composition of both soluble and insoluble cyanophycin produced by *Tm*CphA1 in *E. coli* was next characterized. Here, purified CphB from *Synechocystis* sp. PCC6803 [29] was first used to digest both soluble and insoluble cyanophycin into dipeptide subunits, which were then solubilized and analyzed by mass spectrometry (MS) (Table 1). For insoluble cyanophycin produced by *Tm*CphA1, MS confirmed the presence of a high ratio of arginine relative to lysine, with 93.9% β-Asp-Arg dipeptides and 6.1% β-Asp-Lys dipeptides. This level of arginine in the insoluble fraction is similar to what has previously been reported in studies of other CphA1 enzymes in *E. coli*, which generally range from 90-97% of the total side chain population [8, 30]. In contrast, the soluble fraction produced by *Tm*CphA1 showed a much higher lysine content, with the β-Asp-Lys dipeptide here comprising 35.4% of the total sample. An increase in lysine content was expected as previous studies have shown that higher lysine content enhances the solubility of cyanophycin at near neutral (i.e., intracellular) pH [8].

**Table 1.**
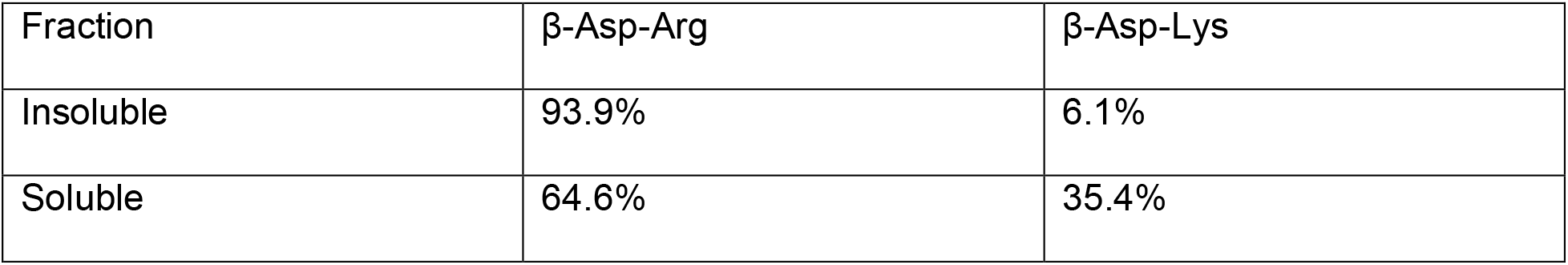
Relative composition of *Tm*CphA1 produced cyanophycin determined by mass spectrometry

Finally, to test if improved pH control and aeration (i.e., relative to the flask scale, which we hypothesized would help to avoid potential ATP limitations) would lead to increased cyanophycin production by *E. coli* expressing *Tm*CphA1, the performance of this strain was investigated in a 1 L bioreactor. Under the conditions examined, dissolved oxygen levels never dropped below 80% and, after 48 hours the OD_600_ within the reactor reached 16.4. At this point, cells were harvested and lysed to recover both soluble and insoluble cyanophycin fractions. A total of 1.80 g of insoluble cyanophycin and 0.12 g of soluble cyanophycin were recovered, which represented a significant improvement over the 1.14 g of insoluble cyanophycin and 0.18 g of soluble cyanophycin produced by the same strain in a 1 L shake flask (where growth reached an OD_600_ of only 11.8). Furthermore, the yields of both insoluble and total cyanophycin from *Tm*CphA1 exceed that which has been reported for other, more commonly studied variants during bench-scale bioreactor production [15]. For comparison, when we applied the same bioreactor conditions for cyanophycin production by *E. coli* harboring *6308*CphA1, only 0.20 g of insoluble material was recovered. Collectively, the outcomes of this study demonstrate that *Tm*CphA1 is a promising CphA1 capable of supporting high level recombinant biosynthesis of cyanophycin in *E. coli*.

## Conclusion

The expression of more stable CphA1 variants was found to support improved yields of insoluble cyanophycin relative to *6308*CphA1 and, in the case of *Tm*CphA1, greater total cyanophycin production. Specifically, relative to *6308*CphA1, *Tm*CphA1 enabled production of 10.8-times and 2.6-times more insoluble and total cyanophycin, respectively. Using a bench-scale bioreactor, cyanophycin production by *E. coli* with *Tm*CphA1 reached as high as 1.92 g/L, including 1.80 g/L of insoluble cyanophycin. Future studies might focus on further production improvements by operating the bioreactor under fed-batch conditions. Overall, with its superior robust and stable expression levels, *Tm*CphA1 possesses strong potential for improving cyanophycin production by engineered *E. coli*. When combined with future strain and process engineering efforts, the outcomes of this study support the robust production of biologically-derived polymers.

## Materials and Methods

### Heterologous cyanophycin production in *E. coli*

T7 driven expression of CphA1 from each of *Tatumella morbirosei* DSM 23827 (*Tm*CphA1), *Acinetobacter baylyi* DSM587 (*Ab*CphA1), and *Synechocystis* sp. UTEX2470 (*Su*CphA1) was achieved using pJ411-derived expression vectors [3]. Because an analogous design constructed for CphA1 from *Synechocystis* sp. PCC 6308 (*6308*CphA1) showed poor expression in preliminary experiments, we instead made use of a previously constructed vector wherein *6308*CphA1 was cloned into pMMB206 and expressed via a tac promoter [18]. All expression vectors were transformed into *E. coli* BL21(DE3), and these strains were used for all expression and cyanophycin production experiments.

Cyanophycin production from the different CphA1s was carried out in 25 mL of terrific broth (TB) inside of 125 mL baffled shake flasks. TB was supplemented with 50 mg/mL kanamycin for *Ab*CphA1, *Su*CphA1, and *Tm*CphA1, and with 35 mg/mL chloramphenicol for *6308*CphA1. Cultures were grown at 37 °C to an OD600 of approximately 0.6, followed by induction of CphA1 expression by the addition of 1 mM IPTG. Following induction, the cultures were shifted to 30 °C for 48 hours for cyanophycin production. Cells were then harvested, and cell pellets stored at −20 °C for further processing.

### Cyanophycin purification

Cell pellets from cyanophycin production were resuspended in 1 M HCl at a ratio of 10 mL of HCl per gram of wet cell weight. Pellets were briefly vortexed and then incubated in a shaker for six hours at room temperature. Samples were then centrifuged at 10,000 g for 15 minutes to remove cell debris, and the supernatant was collected. To help buffer the resulting solution, Tris-HCl pH 8.0 was added to each sample to a final concentration of 0.5 M. This was followed by neutralizing the sample by adding 10 M NaOH. At this stage, the insoluble fraction precipitated and was collected by centrifugation at 10,000 g for 15 minutes. The supernatant, which contains the soluble cyanophycin, was added to twice the volume of ethanol, which causes the soluble cyanophycin fraction to precipitate. Soluble cyanophycin was harvested by centrifugation at 10,000 g for 15 minutes. The insoluble and soluble cyanophycin fractions were lyophilized, weighed, and stored for further analysis.

### Expression and purification of *Tm*CphA1

*Tm*CphA1 was overexpressed using the same strain as for cyanophycin production described above. Cells were grown in LB media supplemented with 50 ug/mL kanamycin at 37 °C. Expression of *Tm*CphA1 was induced at OD600 of 0.6 by the addition of 1 mM IPTG, and protein expression was carried out at 30 °C for 20 hours. Cells were harvested, resuspended in buffer A (150 mM NaCl, 0.5 mM TCEP, 50 mM Tris-HCl, pH 8.0) and lysed by sonication. The lysed sample was centrifuged at 10,000 g for 15 minutes and the supernatant was flown through a 5 mL Ni-NTA column along with 2% buffer B (150 mM NaCl, 0.5 mM TCEP, 500 mM imidazole, 50 mM Tris-HCl, pH 8.0). The Ni-NTA column was then washed with 80% buffer A, 20% buffer B, followed by elution with 100% buffer B. The peak fraction was collected and further purified by SEC on a Superose 6 column equilibrated in buffer A. The SEC peak fraction corresponding to approximately 400 kDa was collected for further analysis.

### *Tm*CphA1 Activity Assay

The assay was performed as described previously [3], with minor modification. A reaction mixture containing 12.5 mM ATP, 2 mM MgCl_2_, 12.5 mM L-aspartic acid, 12.5 mM L-arginine, 12.5 mM KCl, 100 mM Tris-HCl pH 8.0, and a primer of 1.5 mg/mL purified cyanophycin was added to purified *Tm*CphA1 (100 μg/mL final). This reaction mixture with *Tm*CphA1 was incubated for 2 hours, followed by analysis by SDS-PAGE.

### Bioreactor scale cyanophycin production

A 1L Sartorious Biostat A Plus (Sartorious, Göttingen, Germany) bioreactor, was used to test *Tm*CphA1 scale-up. Seed cultures (20 mL) were grown overnight at 37 °C in LB supplemented with 50 μg/L kanamycin. Seed cultures were added to the bioreactor with 1 L of TB, supplemented with kanamycin, and grown at 37 °C with 1 vessel volume per minute (vvm) air sparging, 500 rpm agitation, and held at pH 7.0 until the OD reached 0.5. The pH was controlled through the addition of 6 M NaOH or 6 M HCl. After reaching an OD of 0.5, cultures were induced with 1 mM IPTG, the temperature was reduced to 30 °C, and cultures were left to grow for 48 hours with all other conditions remaining constant. After 48 hours, cells were harvested and cyanophycin was processed as described above.

### Mass Spectrometry

Insoluble and soluble cyanophycin were each resuspended to 25 mg/mL in 50 mM NH_4_CO_3_ and 15 μM cyanophycinase (CphB) and samples were digested overnight at room temperature with mild shaking. Following digestion, the samples were heated to 90 °C for five minutes and then centrifuged at 20,000 g for 5 minutes to remove insoluble material. The samples were then diluted 1:10 in H_2_O and data was collected using a Bruker amaZon speed ETD ion trap mass spectrometer operating at positive ionization mode. Peak areas (including Na^+^ adducts) were used to calculate the relative abundance of β-Asp-Arg and β-Asp-Lys dipeptides.

## References

[1] N. Raddadi, F. Fava, Biodegradation of oil-based plastics in the environment: Existing knowledge and needs of research and innovation, Science of The Total Environment, 679 (2019) 148–158.

[2] K. Ziegler, A. Diener, C. Herpin, R. Richter, R. Deutzmann, W. Lockau, Molecular characterization of cyanophycin synthetase, the enzyme catalyzing the biosynthesis of the cyanobacterial reserve material multi-L-arginyl-poly-L-aspartate (cyanophycin), Eur J Biochem, 254 (1998) 154–159.

[3] I. Sharon, A.S. Haque, M. Grogg, I. Lahiri, D. Seebach, A.E. Leschziner, D. Hilvert, T.M. Schmeing, Structures and function of the amino acid polymerase cyanophycin synthetase, Nature Chemical Biology, 17 (2021) 1101–1110.

[4] R.D. Simon, Cyanophycin Granules from the Blue-Green Alga Anabaena cylindrica: A Reserve Material Consisting of Copolymers of Aspartic Acid and Arginine, Proc Natl Acad Sci U S A, 68 (1971) 265–267.

[5] B. Watzer, K. Forchhammer, Cyanophycin Synthesis Optimizes Nitrogen Utilization in the Unicellular Cyanobacterium Synechocystis sp. Strain PCC 6803, Appl Environ Microbiol, 84 (2018).

[6] T.J. Mueller, E.A. Welsh, H.B. Pakrasi, C.D. Maranas, Identifying Regulatory Changes to Facilitate Nitrogen Fixation in the Nondiazotroph Synechocystis sp. PCC 6803, ACS Synthetic Biology, 5 (2016) 250–258.

[7] A. Steinle, K. Bergander, A. Steinbüchel, Metabolic engineering of Saccharomyces cerevisiae for production of novel cyanophycins with an extended range of constituent amino acids, Appl Environ Microbiol, 75 (2009) 3437–3446.

[8] L. Wiefel, A. Steinbüchel, Solubility behavior of cyanophycin depending on lysine content, Applied and environmental microbiology, 80 (2014) 1091–1096.

[9] J. Aravind, T. Saranya, G. Sudha, K. Palanisamy, A Mini Review on Cyanophycin: Production, Analysis and Its Applications, 2016, pp. 49–59.

[10] J. Du, L. Li, S. Zhou, Microbial production of cyanophycin: From enzymes to biopolymers, Biotechnology Advances, 37 (2019) 107400.

[11] J. Du, L. Li, S. Zhou, Enhanced cyanophycin production by Escherichia coli overexpressing the heterologous cphA gene from a deep sea metagenomic library, J Biosci Bioeng, 123 (2017) 239–244.

[12] C.L. Brady, S.N. Venter, I. Cleenwerck, K. Vandemeulebroecke, P. De Vos, T.A. Coutinho, Transfer of Pantoea citrea, Pantoea punctata and Pantoea terrea to the genus Tatumella emend. as Tatumella citrea comb. nov., Tatumella punctata comb. nov. and Tatumella terrea comb. nov. and description of Tatumella morbirosei sp. nov, Int J Syst Evol Microbiol, 60 (2010) 484–494.

[13] J. van Heijenoort, Recent advances in the formation of the bacterial peptidoglycan monomer unit, Nat Prod Rep, 18 (2001) 503–519.

[14] C.A. Smith, Structure, function and dynamics in the mur family of bacterial cell wall ligases, J Mol Biol, 362 (2006) 640–655.

[15] N. Kwiatos, A. Steinbüchel, Cyanophycin Modifications—Widening the Application Potential, Frontiers in Bioengineering and Biotechnology, 9 (2021).

[16] I. Sharon, S. Pinus, M. Grogg, N. Moitessier, D. Hilvert, T. Schmeing, A cryptic third active site in cyanophycin synthetase creates primers for polymerization, Nature Communications, 13 (2022) 3923.

[17] J. Kroll, S. Klinter, A. Steinbüchel, A novel plasmid addiction system for large-scale production of cyanophycin in Escherichia coli using mineral salts medium, Applied Microbiology and Biotechnology, 89 (2011) 593–604.

[18] N.A. Khlystov, W.Y. Chan, A.M. Kunjapur, W. Shi, K.L.J. Prather, B.D. Olsen, Material properties of the cyanobacterial reserve polymer multi-l-arginyl-poly-l-aspartate (cyanophycin), Polymer, 109 (2017) 238–245.

[19] E. Aboulmagd, F.B. Oppermann-Sanio, A. Steinbuchel, Purification of Synechocystis sp. strain PCC6308 cyanophycin synthetase and its characterization with respect to substrate and primer specificity, Appl Environ Microbiol, 67 (2001) 2176–2182.

[20] E. Aboulmagd, F.B. Oppermann-Sanio, A. Steinbüchel, Molecular characterization of the cyanophycin synthetase from Synechocystis sp. strain PCC6308, Arch Microbiol, 174 (2000) 297–306.

[21] M.V. Merritt, S.S. Sid, L. Mesh, M.M. Allen, Variations in the amino acid composition of cyanophycin in the cyanobacterium Synechocystis sp. PCC 6308 as a function of growth conditions, Arch Microbiol, 162 (1994)158–166.

[22] T. Miyakawa, J. Yang, M. Kawasaki, N. Adachi, A. Fujii, Y. Miyauchi, T. Muramatsu, T. Moriya, T. Senda, M. Tanokura, Structural bases for aspartate recognition and polymerization efficiency of cyanobacterial cyanophycin synthetase, Nature Communications, 13 (2022) 5097.

[23] I. Sharon, S. Pinus, M. Grogg, N. Moitessier, D. Hilvert, T.M. Schmeing, A cryptic third active site in cyanophycin synthetase creates primers for polymerization, Nature Communications, 13 (2022) 3923.

[24] R.D. Simon, The biosynthesis of multi-L-arginyl-poly(L-aspartic acid) in the filamentous cyanobacterium Anabaena cylindrica, Biochim Biophys Acta, 422 (1976) 407–418.

[25] T. Hai, K.M. Frey, A. Steinbüchel, Engineered cyanophycin synthetase (CphA) from Nostoc ellipsosporum confers enhanced CphA activity and cyanophycin accumulation to Escherichia coli, Appl Environ Microbiol, 72 (2006) 7652–7660.

[26] M. Krehenbrink, A. Steinbüchel, Partial purification and characterization of a non-cyanobacterial cyanophycin synthetase from Acinetobacter calcoaceticus strain ADP1 with regard to substrate specificity, substrate affinity and binding to cyanophycin, Microbiology, 150 (2004) 2599–2608.

[27] T. Hai, F.B. Oppermann-Sanio, A. Steinbüchel, Molecular characterization of a thermostable cyanophycin synthetase from the thermophilic cyanobacterium Synechococcus sp. strain MA19 and in vitro synthesis of cyanophycin and related polyamides, Appl Environ Microbiol, 68 (2002) 93–101.

[28] K.M. Frey, F.B. Oppermann-Sanio, H. Schmidt, A. Steinbüchel, Technical-scale production of cyanophycin with recombinant strains of Escherichia coli, Appl Environ Microbiol, 68 (2002) 3377–3384.

[29] I. Sharon, M. Grogg, D. Hilvert, T.M. Schmeing, The structure of cyanophycinase in complex with a cyanophycin degradation intermediate, Biochimica et Biophysica Acta (BBA) - General Subjects, 1866 (2022) 130217.

[30] M. Frommeyer, L. Wiefel, A. Steinbüchel, Features of the biotechnologically relevant polyamide family “cyanophycins” and their biosynthesis in prokaryotes and eukaryotes, Crit Rev Biotechnol, 36 (2016) 153–164.

